# Predicting peptide presentation by major histocompatibility complex class I using one million peptides

**DOI:** 10.1101/349282

**Authors:** Kevin Michael Boehm, Bhavneet Bhinder, Vijay Joseph Raja, Noah Dephoure, Olivier Elemento

**Affiliations:** Caryl and Israel Englander Institute for Precision Medicine, Weill Cornell Medical College, New York, New York, USA; Institute for Computational Biology, Weill Cornell Medical College, New York, New York, USA; Department of Biochemistry, Weill Cornell Medical College, New York, New York, USA; Meyer Cancer Center, Weill Cornell Medical College, New York, New York, USA

## Abstract

Improved computational tools are needed to prioritize putative neoantigens within immunotherapy pipelines for cancer treatment. Herein, we assemble a database of over one million human peptides presented by major histocompatibility complex class I (MHC-I), the largest known database of its type. We use these data to train a random forest classifier (ForestMHC) to predict likelihood of MHC-I presentation. The information content of features mirrors the canonical importance of positions two and nine in determining likelihood of binding. Our random forest-based method outperforms NetMHC and NetMHCpan on test sets, and it outperforms both these methods and MixMHCpred on new mass spectrometry data from an ovarian carcinoma sample. Furthermore, the random forest scores correlate monotonically with peptide binding affinities, when known. Finally, we examine the effect size of gene expression on peptide presentation and find a moderately strong relationship. The ForestMHC method is a promising modality to prioritize neoantigens for experimental testing in immunotherapy.

## Author summary

Neoantigens presented by major histocompatibility complex class I (MHC-I) are critical to modern cancer immunotherapy strategies, including vaccine design, identification of likely responders for checkpoint inhibitors, and CAR-T cell therapies. To act as a neoantigen, a peptide must be presented by at least one of the patient’s MHC-I alleles. Machine learning classifiers exist to predict this presentation, but increasingly available mass spectrometry (MS) data and underexplored machine learning methods offer possibilities for improvement. Herein, we establish the largest known MS database of peptides bound to MHC-I and use it to train improved classifiers. We optimize the selection of features and examine the information content in each feature. Next, we validate the predictor on original data generated for this work, and we compare the predicted scores for peptides with chemical affinity data. Finally, we mine our large database to investigate the effect size of gene expression on peptide presentation.

## Introduction

Immunotherapy targeting neoantigens is uniquely poised to effect precise, powerful treatment of cancer. Among the most promising modalities are adoptive cell transfer (ACT), peptide vaccines, and checkpoint blockade. Specifically, ACT, such as expanded tumor-infiltrating lymphocytes (TILs), chimeric antigen receptor (CAR)-modified T cells, and T cell receptor (TCR)-modified T cells, has demonstrated remarkable clinical efficacy. For example, cellular therapies targeting CD19 have initiated durable complete responses (CR) in patients with follicular lymphoma, chronic lymphocytic leukemia, and other B-cell malignancies [1]. However, these therapies also carry substantial risk. Lethal reactions, including cytokine release syndrome, encephalopathy, and lymphohistiocytosis are strongly associated with the use of CAR-T cells [2]. Furthermore, on-target, off-tumor side effects occur when the peptides targeted are not private to the tumor. In cellular therapy for melanoma, for example, investigators targeted MART-1 and gp100 and noted significant toxicity against the skin, eyes, and inner ear [3]. In other cellular therapies targeting HER2 and MAGE-A3, patients experienced swift and sometimes lethal cardiopulmonary toxicity attributable to titin cross-reactivity [4]. Hence, clinical utility of immunotherapy targeting public antigens is limited by on-target, extra-tumoral action.

Neoantigens are peptides derived from private mutations within the tumor: they are attractive targets because on-target action is limited to the tumor itself. These neoantigens are not present during thymic selection, and thus lymphocytes targeting these epitopes are possibly present within the endogenous pool of lymphocytes [1]. This offers the possibility of reduced costs and complexity compared with engineered lymphocytes. Furthermore, neoantigens are more likely to be derived from mutations critical to oncogenesis: when the neoantigens do derive from driver mutations, this reduces the theoretical risk of antigen escape compared to non-neoantigens [5]. Preliminary clinical evidence is promising: one patient with cholangiocarcinoma experienced a partial response lasting at least two years after infusion of TILs specific to a mutation in her tumor [6]. Beyond targets for cellular infusions, neoantigens also are predictive of response to checkpoint blockade: mutational burden in melanoma correlates with clinical response to CTLA-4 inhibitors [7]. Clinically, one study demonstrated the presence of TILs specific to a neoantigen in a patient’s tumor, and immunologic response of those cells rose during treatment with ipilimumab [8]. Neoantigens also permit targeted peptide-based vaccines. In mice, injections of neoantigens caused comparable clinical response to checkpoint blockade [9]. Despite this promising preliminary evidence, identifying neoantigens consistently for these therapies remains elusive and complex.

Toward clinical application of neoantigen-based therapy, the exome of the tumor is first sequenced and compared to the exome of germline cells (typically the blood), yielding a set of private mutations within the tumor. These mutations are converted into putative neoantigens, peptides of approximately 8-12 amino acids (most typically length 9). This list of putative neoantigens is too extensive for exhaustive testing for clinical therapy. One approach to determine which few of these putative neoantigens truly are presented and possibly immunogenic in the tumor, is to use mass spectrometry (MS) to examine the entire tumoral immunopeptidome. This method, though highly accurate and thorough, is costly and time-intensive. Furthermore, it requires a relatively large amount of sample from the patient (up to 1cm^3^), which cannot always be obtained [10]. Thus, a more efficient method is needed to guide therapy.

*In silico* prediction is another method to guide neoantigen-based therapies. This technique takes as input a list of the potential peptides derived from private mutations and prioritizes them by likelihood of presentation. Of course, prediction must be tailored to the patient’s specific alleles of human leukocyte antigen (HLA) A, B, and C, which code for MHC-I. Each variant of MHC-I has a distinct preference for a binding motif: hence, the specific alleles determine the space of possible peptides presented within the tumor. Multiple predictors are publically available to rank potential neoantigens.

One widely used approach is NetMHC [11,12]. This method uses artificial neural networks (ANN) as its underlying machine learning framework. For training data, NetMHC uses affinity data measured from *in vitro* assays. A related predictor, NetMHCstabpan, uses the half life of the MHC-peptide complex *in vitro* [13]. However, these data limit the application of NetMHC and NetMHCstabpan to neoantigen prioritization: actual presentation *in vivo* is contingent on other processes unrelated to chemical affinity. These include proteasomal processing, abundance of proteins containing specific sequences, and biological half-life [14,15]. Hence, NetMHC and NetMHCstabpan are suboptimal for neoantigen prioritization because of their reliance on chemical training data.

Increasingly available MS datasets are more suitable for training predictors of peptide binding: because these data describe epitopes actually presented *in vivo*, they account both for chemical affinity and biological processes required for presentation. Furthermore, sampling the immunopeptidome does not require *a priori* peptide synthesis or selection, and this reduced bias increases the theoretical likelihood of discovering novel motifs and true binders [14]. Two publically available methods make use of MS datasets: NetMHCpan is an ANN-based predictor trained on both affinity data and MS data [16]. MixMHCpred is trained only on MS data, and it employs position weight matrices (PWMs) established by a mixture model for each allele [10].

Though these methods are trained on appropriate MS data, there has been insufficient exploration of alternative features and machine learning frameworks. Indeed, the developers of MixMHCpred did not attempt to optimize the machine learning framework of the method [10]. Though NetMHCpan uses ANN, it relies upon BLOSUM encoding as input features, and additional biochemical features and sequence representations have the potential to improve performance [16]. Herein, we investigate various features and machine learning methods trained on public MS data to optimize predictors of peptide binding.

## Results

### Database characteristics

The total number of peptides collected from the Proteomics Identifications Database (PRIDE), SysteMHC Atlas, and other published data (see methods) was 1.03E6. To our knowledge, this is the largest database of its type to date. Of these peptides, 5.7E5 (55%) are nine amino acids in length (**Fig S1**). MHC-I alleles tend to have a strong preference for peptides of length nine, making this length the priority for classification. Of the nonamers, 2.9E5 (51%) were reported in polyallelic samples. We deconvoluted these peptides using MixMHCpred, with 2.8E4 peptides discarded due to unavailable predictions for the given alleles and 4.3E4 peptides discarded due to a confidence in allele assignment of less than 95%. We then pooled the peptides by allele, merging the deconvoluted peptides with the peptides from monoallelic sets and from datasets already presented as deconvoluted using NetMHC. During this pooling, we included only unique peptides (3.3E5 peptides were duplicates). The total number of unique nonamers assigned to alleles was 1.6E5.

The cell lines in the database spanned B-cell lymphoblasts, breast cancer, leukemia, lymphoma, glioblastoma, melanoma, fibroblasts, embryonic kidney cells, and colon carcinoma. The clinical samples included peripheral blood mononuclear cells, melanoma, meningioma, and lung cancer. The number of MHC-I alleles was 82, including 65 alleles resolved to four digits and 17 alleles resolved to two digits. We had 26 HLA-A alleles, 40 HLA-B alleles, and 16 HLA-C alleles.

### Feature selection

We began by finding the optimal combination of features considered. Namely, we considered hydropathy, blosum62 sequence encoding, one-hot (sparse) sequence encoding, presence of an aromatic residue, mass, and charge at physiological pH. We chose these features by a combined review of biochemical and MHC-I binding predictor literature [10,12,17]. In particular, we chose the aromatic feature due to experimental evidence of allosteric networks regulating the conformation of MHC-I binding grooves in a selected allele [10].

To identify the optimal feature combinations, we built random forest classifiers for all 82 alleles across 63 possible feature subsets, sizes one to six. For the training set, we used a 1:1 ratio of randomly generated nonamers from SwissProt to true binders. For the test set, we used a 99:1 ratio of these random decoys to true binders. We employed precision in the top 1% of predictions (Prec1%) and area under the precision recall curve (AUPRC) to measure performance, and we calculated mean performance across 82 alleles on the test set (**Fig 1**).

**Fig 1.**
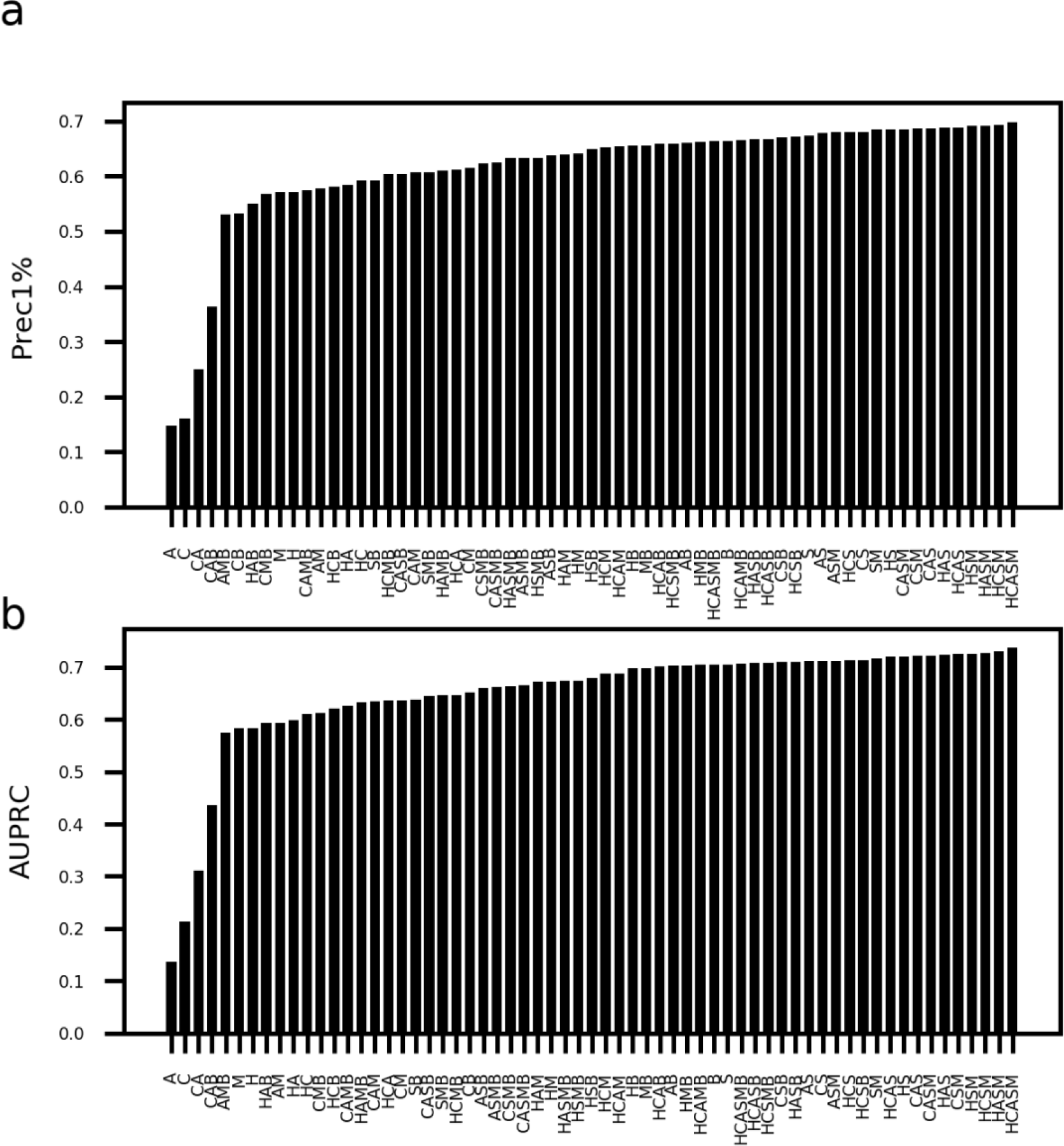
Performance across all combinations of investigated features. We compared performance across 63 feature subsets for all alleles, with (a) showing Prec1% and (b) showing AUPRC for each feature combination. (H- hydropathy, A- presence of aromatic, C- charge at physiological pH, M- mass, S- sparse encoding, B- blosum62 encoding)

A number of feature sets yielded excellent performance, with sparse encoding present in all ten of the top ten feature sets for both AUPRC and Prec1%, and with blosum62 encoding present in 0/10 for both. Hydropathy was present in 7/10 for both, and mass, charge, and aromaticity were present in 6/10 for both. Based on this analysis, we chose the combination of hydropathy, presence of aromatic rings, sparse encoding, and mass (HASM): this combination yields performance within the top 1% of maximal Prec1% values and AUPRC values, and it has one fewer feature than the top performer (HCASM), reducing the likelihood of overfitting. The final random forest classifiers use the HASM feature combination.

### Comparison to existing predictors on test data

Using the combination of hydropathy, presence of an aromatic ring, sparse encoding, and mass features, we trained a random forest model for each allele. For these models, we used 1000 trees, gini impurity, and the square root of the total number of features as a maximum. Decoy peptides of length nine were again generated randomly from SwissProt for a 1:1 class balance during training and 99:1 class balance during testing.

Our final set of random forest (RF) classifiers achieved an average Prec1% of 0.69 and AUPRC of 0.73 across test sets by five-fold cross validation. We compared the performance of our RF classifiers to other publically available classifiers—NetMHC (Prec1% 0.54, AUPRC 0.51), NetMHCpan (Prec1% 0.64, AUPRC 0.65), NetMHCstabpan (Prec1% 0.46, AUPRC 0.41), and MixMHCpred (Prec1% 0.70, AUPRC 0.74). The results across all alleles by five-fold cross-validation on the test sets are shown in **Fig 2**. By the Mann-Whitney U Test, our RF-based method outperformed NetMHC, NetMHCpan, and NetMHCstabpan. There was no significant difference between the RF method and MixMHCpred.

**Fig 2.**
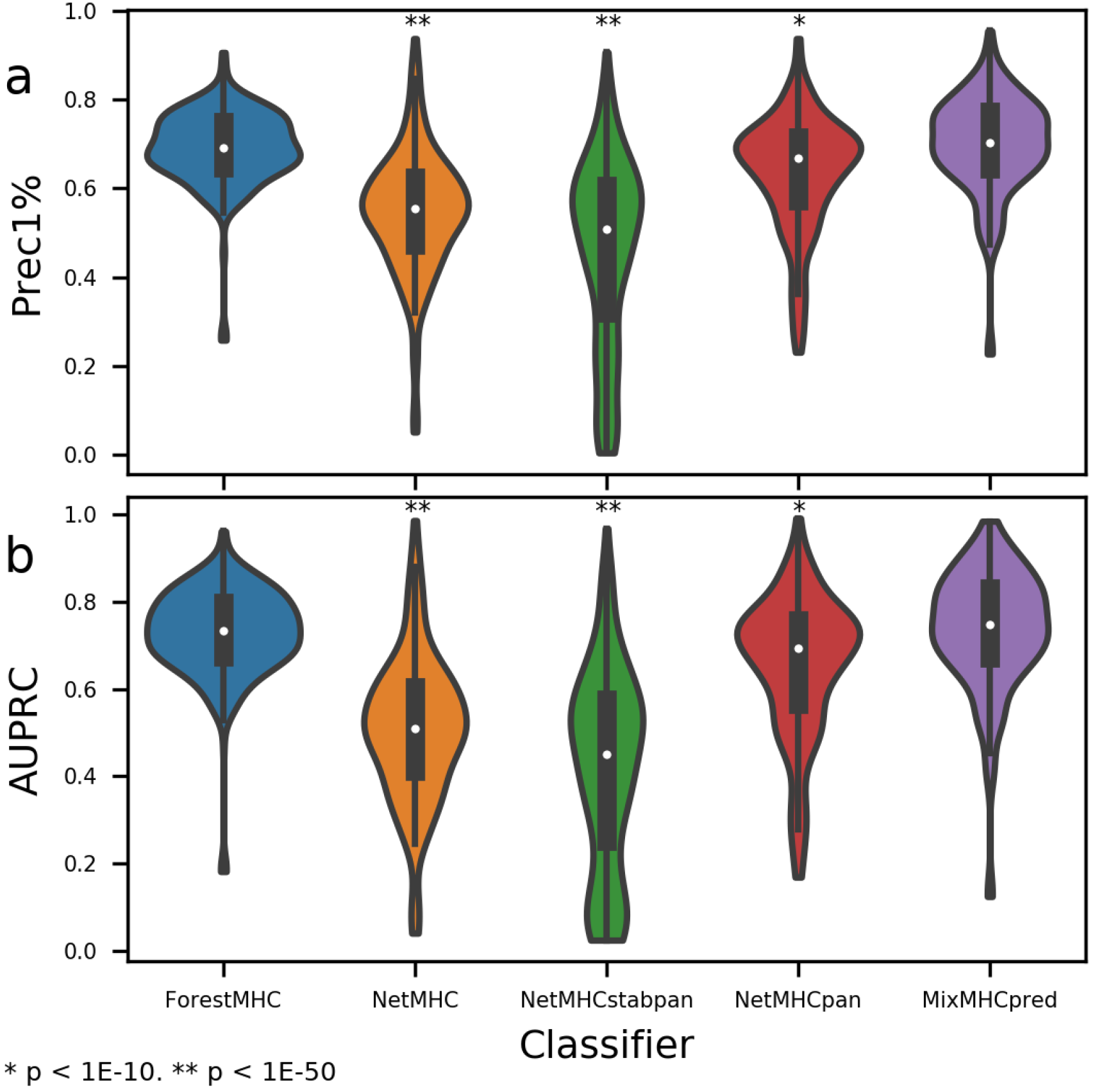
RF-based method outperforms existing predictors on unbalanced data. (a)AUPRC and (b) Prec1% are greater for RF compared to NetMHC, NetMHCpan, and NetMHCstabpan, with no significant difference between MixMHCpred and RF (p > 0.01). P values are by Mann-Whitney U Test compared to ForestMHC.

It was expected that, by this methodology of testing, the performance of our method could not exceed that of MixMHCpred for two reasons. First, many of the data in our database also were used to train MixMHCpred (MMP). Hence, some peptides assigned to our test set (drawn at random from the data) were likely included in the training set during the development of MMP. Second, we relied upon MMP to deconvolute 51% of our peptides, and we discarded all peptides without available MMP predictions or with a confidence of less than 95% in the assignment. Thus, the test dataset is biased in favor of high-certainty peptides for MMP and also contains peptides included in the training of MMP. Given these conditions, it is remarkable that this new method performs at a level that is statistically indistinguishable from MMP.

### Feature importance analysis

We next wondered about the information content of each feature. To measure this, we calculated the mean reduction in Gini impurity at nodes using each feature across all trees in each ensemble. We then averaged this quantity arithmetically across all classifiers (**Fig 3**). Information is higher, on average, in the hydropathy and mass features than the sparse encoding. Positions two and nine contain substantially more average information within the hydropathy, aromaticity, and mass features, and the information for one-hot encoding is higher for positions two and nine compared to other positions. To rule out the potentially confounding influence of deconvolution, we repeated the analysis using only mono-allelic data: findings were similar (**Fig S2**). The importance of these features is corroborated biologically: most known MHC-I alleles prefers characteristic amino acids found at two and nine, the canonical anchor residues [18].

**Fig 3.**
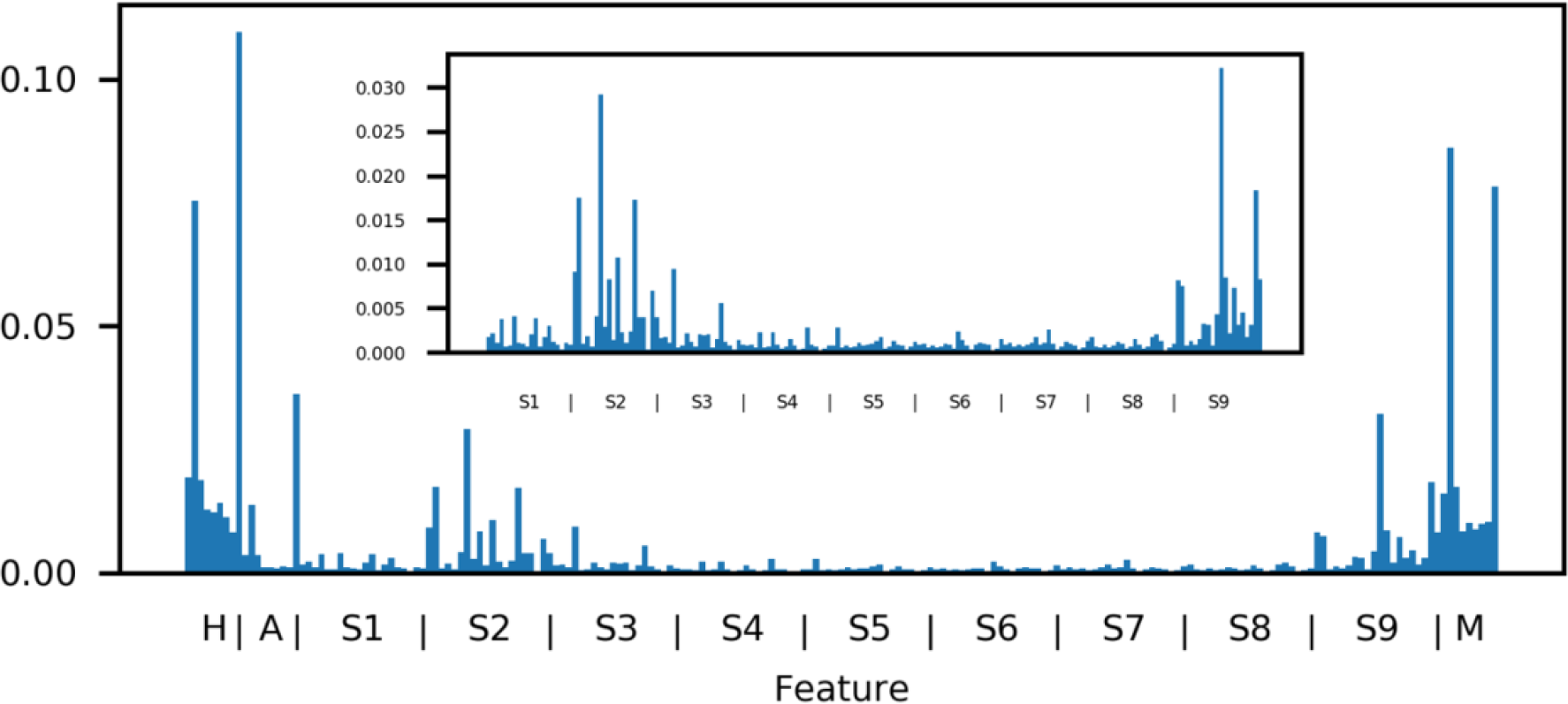
Mean information by feature mirrors canonical anchor residues. The mean information (reduction in Gini impurity) across all classifiers is shown, with the inset showing only the feature subset of sparse encoding. Note that the information is higher at positions two and nine for the sparse encoding features (each a 20-dimensional encoding of each amino acid). For the nine-dimensional features of hydropathy, aromaticity, and mass, the mean information content at positions two and nine is substantially higher than at other positions.

### Validation on never-before-seen data

A more rigorous, realistic test is the application of classifiers to data that is both new (never seen by any classifier) and polyallelic (requiring ranking while taking into account multiple alleles). We performed an experiment to elute ligands bound to MHC-I in an ovarian carcinoma cell line (SK-OV-3), identify them using mass spectrometry (see Methods). We obtained 694 high-confidence peptides. We mixed the 534 resultant nonamers computationally with a 99-fold excess of random decoys. To each classifier, we provided the HLA alleles (obtained from Adams et al.) and the list of mixed true peptides and random decoys [19]. Prec1%—calculated with five different sets of decoys mixed in—was higher than all other methods tested (**Fig 4**). Our classifier outperformed MixMHCpred, NetMHC, NetMHCstabpan, and NetMHCpan. These results demonstrate the promise of RF and these features to supercede existing methods of epitope prioritization.

**Fig 4.**
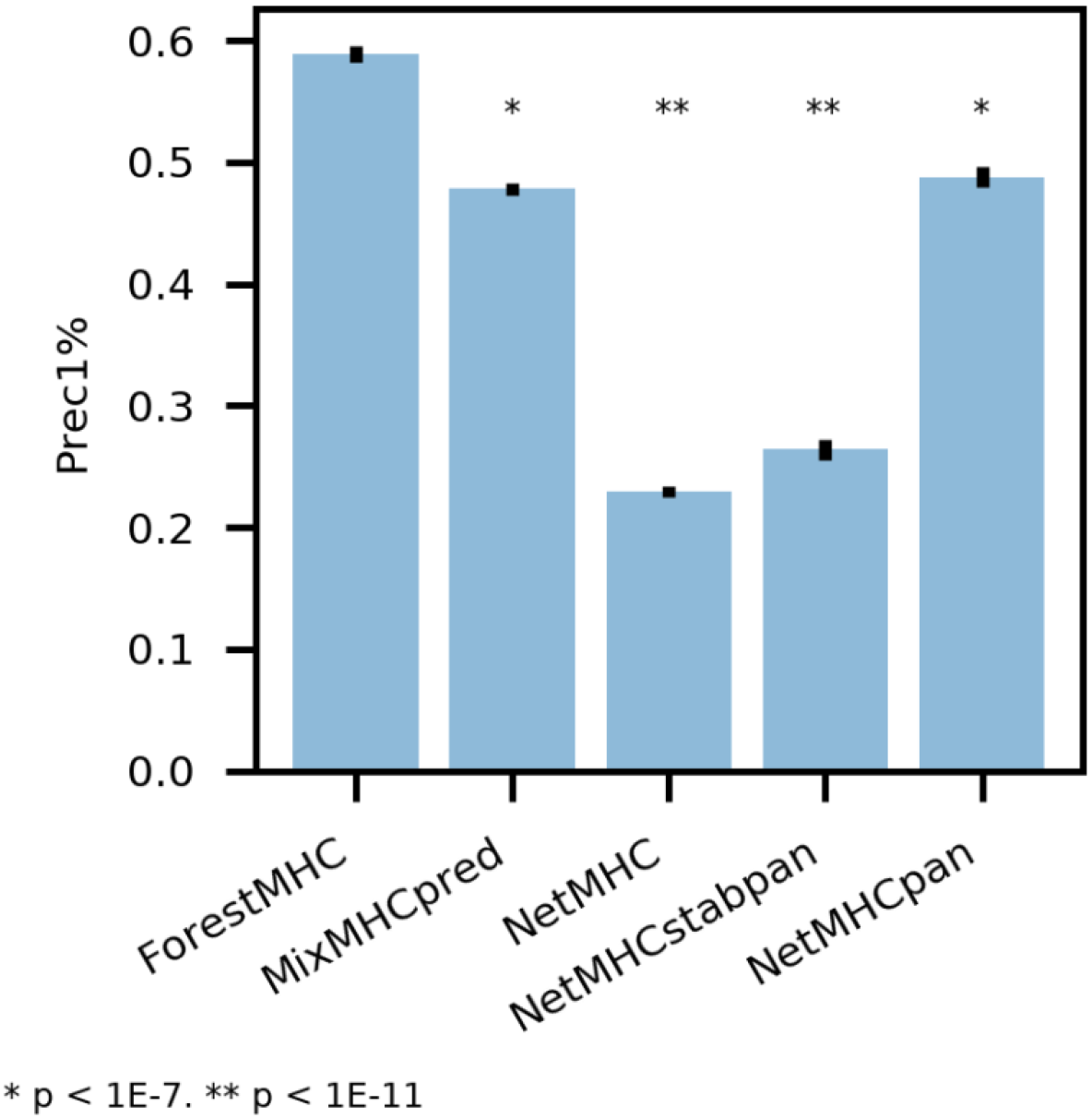
New method outperforms existing classifiers on never-before-seen MS data. Precision in the top 1% of predictions for our method is superior compared to Prec1% of existing methods on newly generated data from ovarian carcinoma cell culture. P values are by Student’s t-test.

### Comparison with other machine learning methods

We evaluated several other methods of machine learning, including deep artificial neural networks, but we consistently noted lower performance than random forests (**Fig S3**). All classifiers were trained on the same database and tested on our new data from ovarian carcinoma cells with 99-fold excess of random decoys, using the established four feature sets (HASM). ForestMHC consistently performed better, with a mean Prec1% of 0.59 across five different sets of random decoys. Deep neural networks (mean Prec1% 0.41), convolutional neural networks (mean Prec1% 0.34), and support vector machines (mean Prec1% 0.07) did not perform as well. The especially low performance of the SVM is expected given the importance of nonlinear interactions among residues in establishing the specificity for binding by MHC-I. Furthermore, RF’s outperformance of convolutional and deep neural networks—other nonlinear methods—demonstrates the potential utility of this machine learning technique in peptide binding classification.

### Correlation of RF score with affinity

We next wondered how the RF scores related to experimentally measured affinity data. Using all nonamers with available IC_50_ data on the Immune Epitope Database, we generated RF scores using our predictors and assessed the correlation with IC_50_ values (**Fig 5**). The relation is of moderately high monotonicity, with a mean Spearman’s coefficient of −0.59 (range: −0.16, −0.79) across 22 alleles, weighted by number of entries. The relationship is weakly linear, with a mean Pearson’s coefficient of −0.27 (range: −0.09, −0.73).

**Fig 5.**
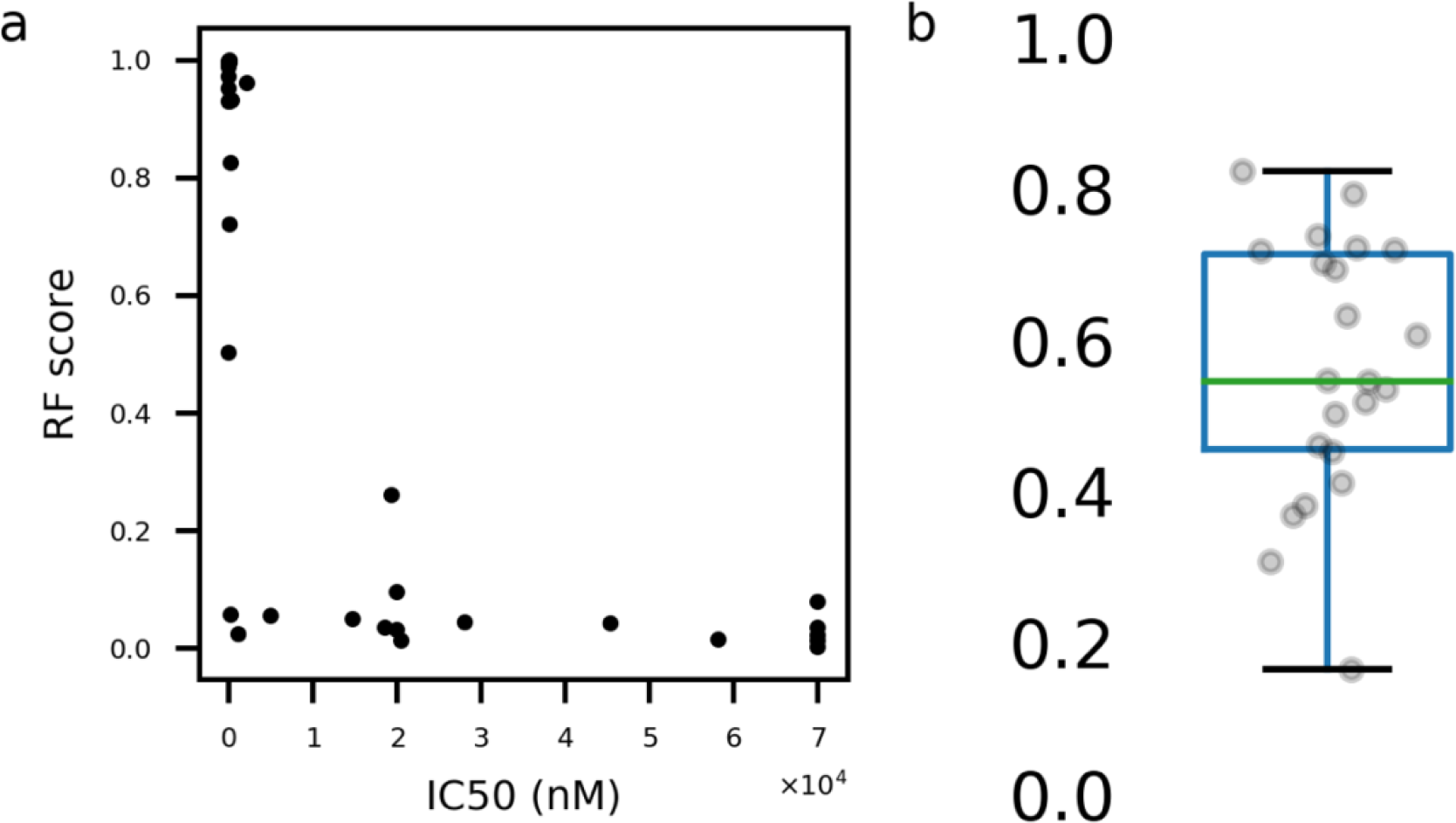
RF score correlates monotonically with IC50 affinity. (a) Example plot shows data from HLA-A29:02. (b) Spearman coefficients for IC50 vs RF score by allele; box plot shown is unweighted and shows IQR within box and median by line within.

This type of relationship—monotonic, but not necessarily linear—is sensible for these two quantities: while the IC_50_ measurements contain only information about ligand binding, MS elution datasets contain information about whether the peptide is actually found bound to MHC-I biologically. The latter process is complex and depends on proteasomal processing and abundance of source proteins, among other factors. Furthermore, chemical affinity data require *a priori* selection of epitopes to test, which limits the space of the immunopeptidome explored [14].

### Effect of gene expression on peptide presentation

Previous studies suggest that peptides derived from proteins coded by highly expressed genes are more likely to be presented by MHC-I [15,20]. Using our large database, we sought to validate this claim. Using mRNA gene expression data for each cell line and clinical sample, or for its closest proxy, we compared the expression of genes that code for peptides presented by MHC-I to those that do not (**Fig 6**). The mean Cliff’s d value was 0.60 (range: 0.47, 0.76) when unweighted and 0.59 when weighted by the number of genes successfully mapped from proteins. Hence, mining our large database corroborates previous findings that gene expression has a large positive effect on presentation by MHC-I. These data may prove predictive as features in future iterations of ForestMHC.

**Fig 6.**
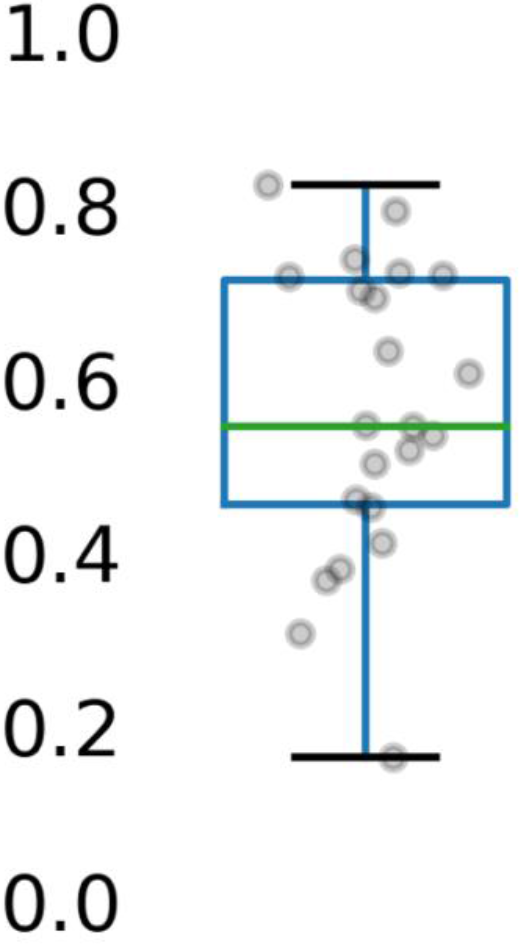
Gene expression has a modestly large positive effect on peptide presentation. The Cliff’s d values shown here are for pooled cell lines and clinical samples within the database

## Discussion

Herein, we have established a new random-forest approach to prioritize putative neoantigens for experimental investigation. Our method outperforms other methods on new data not found in the training set of any classifier, and it is a promising option for future epitope prioritization. The relative feature importance of positions two and nine dovetails well with existing knowledge about anchor positions for MHC-I. The RF scores correlate monotonically with IC50 values, and analysis of our large database corroborates the positive effect of genetic expression on presentation of derived peptides by MHC-I.

One limitation of ForestMHC is its reliance on wild-type peptides rather than neoantigens for training data. Given this constraint, the predictions describe whether a given peptide is likely to bind to an allele of MHC-I, not whether that peptide is also necessarily immunogenic. In the future, we will evaluate ForestMHC’s ability to identify true neoantigen binders. Furthermore, not all peptides presented by MHC-I are of length nine, and future work must also include support for peptides of other lengths. Finally, there are insufficient MS data to train classifiers for the majority of HLA 1 alleles, and we did not train any classifiers to predict binding to HLA 2. As more MS data become available, we will continue to extend this coverage.

## Methods

### Dataset and pre-processing

We acquired publically available mass spectrometry peptide elution datasets from PRIDE (https://www.ebi.ac.uk/pride/archive/) [21,22], SysteMHC (https://systemhcatlas.org/) [23], and supplementary files of individual publications, for a total of 24 distinct data sets [10, 14, 20, 24–41]. Only datasets with false discovery rates of 5% or lower were included. We excluded peptides if their length was not nine amino acids or if they included any amino acids outside of the standard set of twenty. We pooled mono-allelic data by allele and deconvoluted poly-allelic data using MixMHCpred with a p value threshold of 0.05 before pooling [15]. We discarded entries for which MixMHCpred predictions were unavailable, and we also discarded duplicate entries for a given allele. We trained classifiers only for alleles with 50 or more peptides from MS datasets. For class balance during training, we added randomly generated nonamers from SwissProt for a 1:1 ratio (uniprot.org). For testing, the ratio of decoys to true binders was 99:1 ratio.

### Machine learning frameworks

We trained one classifier for each individual allele. For the random forest approach, we used 1000 trees, allowed the square root of the total number of features at each decision node, and performed bootstrapping. For the convolutional neural network approach, we used a modified version of the approach taken by Hu & Liu [42]. We encoded each amino acid in twenty channels representing the standard amino acids. By layer, we convolved this input with 512 filters (kernel size: 2, stride: 1), derived the max pool (kernel size: 2, stride: 2), convolved with 512 filters again (kernel size: 3, stride: 1), flattened, processed by a fully-connected layer (400 units, ReLU activation function), discarded using a dropout layer (40% dropout), and finally fed this result into two logits. We used cross entropy with softmax to calculate loss. For the deep neural network approach, we used two fully connected layers of 500 and 100 units.

### Feature engineering

We chose features from among blosum62 encoding, sparse encoding, hydropathy score, indicator of presence of an aromatic ring, molar mass, and charge of the amino acid at physiological pH. To determine the optimal subset, we conducted an exhaustive search of all possible subsets of sizes from one to six, inclusive. We defined information per feature as reduction in Gini impurity at nodes using each feature (averaged across all trees in the ensemble), and we averaged this quantity across all classifiers.

### Performance metrics

To measure performance of our classifiers, we calculated Prec1% after mixing true binders with a 99-fold excess of random decoys from SwissProt. This metric has been used by others in the development of classifiers, and it is attractive because of its encapsulation of real-world applications for the classifiers [14,15]. That is, the classifiers produced herein are designed for prioritization of putative neoantigens for experimental testing by immunologic assays. The best measure of a useful classifier, thus, is its ability to prioritize truly bound peptides over the noise of random sequences. Hence, we established Prec1% as the principal metric.

As a secondary metric, we chose the AUPRC. Though the AUPRC is less directly translatable to the intended use of these classifiers, the metric also is useful to evaluate the relative proportion of true positives within predicted positives [43]. Furthermore, the AUPRC better reflects a classifier’s ability to separate highly unbalanced datasets compared to the area under the receiver operating characteristic curve (AUROC). While the AUROC has a value of 0.5 for a random classifier no matter the ratio of negatives to positives, the AUPRC’s value for a random classifier is the ratio of the positives to negatives [43]. Hence, with our ratio of cases and controls, the AUPRC value would be 0.01 for a random classifier.

Mean Prec1% and AUPRC were calculated by five-fold stratified cross validation on the test set. The test set consisted of 25% of the known binders to MHC Class I along with a 99-fold excess of decoys from SwissProt. For each iteration of the cross-validation, we used the same test set for all classifiers, namely MixMHCpred 1.1, NetMHCpan 4.0, NetMHC 4.0, and NetMHCstabpan 1.0.

### SK-OV-3 MHC-I Peptide Identification Methods

#### Cell Line and Antibody

We characterized the HLA-1 peptidome of an ovarian carcinoma cell line, SK-OV-3 (ATCC HTB-77). W6/32 monoclonal antibody (Bio X Cell, Catalog #BE0079) was cross-linked to Protein-A Agarose (Santa Cruz sc-2001) beads using dimethyl pimelimidate (D8388 Sigma).

#### Purification of HLA-1 Complexes

We conducted the experiment in accordance with the procedure outlined by Bassani-Sternberg et al. [15]. Briefly, we lysed a single pellet of 3E7 SK-OV-3 cells with 0.25% sodium deoxycholate, 0.2mM iodoacetamide, 1 mM EDTA, 1:200 Protease/Phosphatase inhibitors (Thermo), 1mM PMSF, and 1% octyl-β-D glucopyranoside (Sigma) in PBS at 4°C for one hour. The lysate was cleared for one hour at 20,000 × g prior to immunoaffinity purification of HLA-1 molecules with the cross-linked W6/32 antibody. We then washed beads with 10 × bead volume of 150mM NaCl, 20mM Tris.HCl (buffer A), 10 volumes of 400mM NaCl, 20 mM Tris.HCl, 10 volumes of buffer A again, and lastly with seven volumes of 20mM Tris.HCl, pH 8.0. Next, we eluted HLA-1 molecules by the addition of 500μl of 0.1 N acetic acid at room temperature in two steps following a five-minute incubation each time.

#### Purification and Concentration of HLA-1 Peptides

We loaded HLA complexes and eluted HLA-1 peptides onto a pre-equilibrated Sep-Pak tC18 column (Waters, Milford, MA) and washed with excess 1% formic acid. Bound peptides were eluted with 70% acetonitrile (ACN) and 1% formic acid before being lyophilized.

#### LC-MS/MS Analysis of HLA-1 Peptides

Peptides were reconstituted in 5% formic acid and analyzed by LC-MS/MS on a Thermo Orbitrap Fusion Mass Spectrometer. We separated peptides by reverse-phase HPLC on a hand-packed column (packed with 40 cm of 1.8 μm, 120 Å pores, Sepax GP-C18, Sepax Technologies, Newark, DE) using a 75 minute gradient of 5-25% buffer B (ACN, 0.1% FA) at a 350 nl/min. Peptides were detected using a Top20 method. For each cycle, we acquired one full MS scan of m/z = 375–1400 in the Orbitrap at a resolution of 120,000 at m/z with AGC target = 5×105. Each full scan was followed by the selection of up to 20 of the most intense ions for CID and MS/MS analysis in the linear ion trap. Selected ions were excluded from further analysis for 40s. We also rejected ions with unassigned charge or charge of +1. Maximum ion accumulation times were 100 ms for each full MS scan and 35 ms for MS/MS scans, and all scans were collected in centroid mode.

#### Mass Spectrometry Data Analysis of HLA Peptides

We searched data separately against two different databases using SEQUEST [44]. One search used a set of >200,000 previously identified MHC-I bound peptides downloaded from the Immune Epitope Database (iedb.org) and a null enzyme digestion specificity: that is, only the complete sequences as downloaded were considered as potential matches. A second search used the complete set of reviewed human protein sequences from Uniprot [45], including splice isoforms. This search was performed with “no enzyme” specificity which considers all possible peptide sequences >6 amino acids and <3500 daltons total MH+. We used a composite database containing the translated sequences of all predicted open reading frames of the human genome and their reversed complement to enable target-decoy filtering. We used the following search parameters: a precursor mass tolerance of ±20 ppm, 1.0 Da product ion mass tolerance, no enzyme specificity, a static modification of carbamidomethylation on cysteine (+57.0214), and a dynamic modification of methionine oxidation (+15.9949). We filtered peptide spectral matches to a FDR of 1% using the target-decoy strategy [46] combined with linear discriminant analysis (LDA) using SEQUEST scoring parameters including Xcorr, ΔCn′, precursor mass error, and charge state [47].

### Application of classifiers to SK-OV-3 dataset

We established alleles using data from Adams et al [19]. We mixed the true binders with a 99-fold excess of decoys generated from SwissProt, and then we applied the random forest classifiers for the known HLA alleles. The rank of peptides was determined by the maximum of their random forest scores across all six HLA alleles. We repeated this testing five times, with different sets of random decoys mixed in each time.

### Analysis of effect of gene expression on presentation

We pooled the lists of source genes for presented peptides by cell line or clinical samples across studies. Transcriptomes for given cell lines and samples—or, when unavailable, closely matched proxies—were from NCBI Gene Expression Omnibus and EBI Expression Atlas [48,49]. We used the approach taken by Pearson et al to analyze the effect size of gene expression [20]. Cliff’s d value described the effect size; we included all samples with more than 50 genes successfully mapped from peptides, and we weighted the mean across samples by the number of genes in each sample.

### Correlation of affinity and RF score

From the Immune Epitope Database (IEDB, iedb.org), we downloaded all existing IC50 data for HLA-A, B, and C [50]. We excluded any allele with fewer than 25 entries or for which no random forest classifier was available. For the alleles with sufficient affinity data and a trained classifier, we generated RF scores and correlated them with IC50 values using Spearman’s correlation to evaluate for monotonicity. We calculated the mean coefficient by weighting according to the number of entries for each allele.

## Acknowledgments

This study makes use of data generated by the Blueprint Consortium. A full list of the investigators who contributed to the generation of the data is available from www.blueprint-epigenome.eu. Funding for the project was provided by the European Union’s Seventh Framework Programme (FP7/2007-2013) under grant agreement no 282510 – BLUEPRINT.

